# Investigating the two regimes of fibrin clot lysis: an experimental and computational approach

**DOI:** 10.1101/2021.04.14.439801

**Authors:** Franck Raynaud, Bastien Chopard, Alexandre Rousseau, Karim Zouaoui Boudjeltia, Daniel Monteyne, David Perez-Morga

## Abstract

It has been observed *in vitro* that complete clot lysis is generally preceded by a period of latency during which the degradation seems to be inefficient. However, this latency was merely notified but not yet quantitatively discussed. In our experiments we observed that the lysis ubiquitously occurred in two distinct regimes, a slow and a fast lysis regime. We quantified extensively the duration of these regimes for a wide spectrum of experimental conditions and found that on average the slow regime lasts longer than the fast one, meaning that during most of the process the lysis is ineffective. We proposed a computational model in which the two regimes result from a spatially constrained kinetic of clot lysis: first the biochemical reactions take place at the outer core of the fibrin fibers composing the clot, then in the bulk resulting in the observed fast lysis regime. This simple hypothesis appeared to be sufficient to reproduce with a great accuracy the lysis profiles obtained experimentally. Our results shed light on new insights regarding the dynamical aspects of the lysis of fibrin rich clots in a context where the timing is so critical for patient treatment and outcome.

**Significance:** While the interplay between the main components of the fibrinolytic system is well understood, some dynamical aspects of the fibrinolysis remain unclear. Notably, we observe that *in vitro* fibrin rich clots undergo a slow and inefficient phase of degradation when subject to endogenous fibrinolysis. In fact, it turns out that a large part of the lysis process operates in this slow regime. To explain this observation, we proposed a computational model in which the properties of the binding of the proteins change during the lysis. First plasminogen and tissue plasminogen activator bind at the surface of the fibers, resulting in a slow lysis, then in the bulk of the fibers thus speeding up the degradation of the clot..

## INTRODUCTION

Fibrinolysis is the process by which a thrombus is dissolved. The fibrinolytic system removes the fibrin clot once it has achieved its hemostatic function. By nature, fibrinolysis is delayed, at least relative to coagulation: fibrin is formed in a matter of minutes or even seconds-generally serving a useful purpose, such as the healing of a breach in the vascular wall; afterwards it has to be removed over a period of hour or days while in the meantime tissue repair can take place. Fibrinolysis uses elements from plasma, platelets, tissue, and other blood cells to regulate the degradation of fibrin. This is brought about by the conversion of plasminogen, a plasma protein that circulates as a zymogen to the serine protease, plasmin. Plasmin is a trypsin-like enzyme that cleaves a wide variety of plasma proteins. However, many of these reactions require extremely high levels of plasmin. The physiological function of plasmin is limited primarly to the degradation of the fibrin clot and extracellular matrix molecules. Conversion of plasminogen to plasmin is achieved by a variety of plasminogen activators that exist in tissues throughout the body[1]. The plasminogen activators that are important in this process include tissue-type plasminogen activator and urokinase-type plasminogen activator. Several studies have reported that the risk of ischemic cardiovascular events (CVE) is increased in patients with impaired fibrinolytic function [2–5]. Fibrinolytic activity is primarily determined by the balance between the levels of tissue plasminogen activator (t-PA) and plasminogen activator inhibitor 1 (PAI-1). The endothelial cells are responsible for the production and blood release of t-PA and of PAI-1 to some extent. Multiple factors, such as lipoproteins, cytokines, and inflammatory markers, modulate endothelial cells to produce t-PA and PAI-1[6]. In this work, we focus our attention on acute ischaemic strokes. Indeed, in 2016 the World Health Organization identified stroke as the second causes of death for both sexes and across all ages[7]. Clinical interventions can be classified into either chemical (thrombolysis) or mechanical (thrombectomy) treatment. In particular combination of both appeared to be a promising approach regarding patient outcomes [8, 9]. However the time window for the treatment at the onset of the stroke remains the most critical parameter and patients who benefited from early intervention are associated with significant better 3-month prognosis [10] but treatment success is highly variable [11]. Meanwhile a large effort is encouraging *in silico* trials for thrombectomy and thrombolysis [12], a better understanding of the origin of such variability is still an issue from the clinical perspective and also remains a critical challenge from a modeling point of view. The blood coagulation cascade and fibrinolysis process are a striking example of how the whole is way more complex than its parts. Each component (including fibrin polymerization and fibrinolysis) is well understood if considered individually, isolated form other parts. However taken as a whole, hemodynamics, non-linearities, feedback loops and the variability in physiological conditions increase dramatically the complexity of the coagulation system. Even though it is recognized that computer simulations is an essential tool to deal with this complexity, mathematical modeling remains challenging. Compartmental models describing the biochemical processes of the coagulation and the lysis of the clot require dozens of differential equations [13, 14] as well as advanced sensitivity analysis to decipher the most relevant components of the reactions [15, 16]. In this work we combine in vitro experiments and computational modeling analysis of fibrin rich clot formation and lysis. The dynamics of the processes was monitored from the time of clot formation to the time complete lysis. A computerized semi-automatic device based on turbidity measurements was used to determine the amount of fibrin present in the system. We investigated the effect of changing the concentration of the main components of the fibrinolytic system (fibrinogen, plasminogen, tissue plasminogen activator and Plasminogen activator inhibitor-1) on the clot formation time and the total lysis time. Analysis of turbidity time curve reveals that for all experimental conditions, the lysis of fibrin rich clots is characterized by two regimes: a slow regime where the change in turbidity is linear and a fast regime where the change is exponential. The goal of our computational model is to propose a mechanisms explaining the slow regime observed in our experiments. We tested the hypothesis that the slow (resp. fast) regime corresponded to lysis kinetics occurring at the surface (resp. in the bulk) of the fibers.

## MATERIALS AND METHODS

### Experimental procedures and data acquisition

Our experiment consist of mixing fibrinogen, plasminogen, tPA and PAI-1 in various concentations in a test-tube. Then an excess of thrombin is added and the attenuation of a light beam through the test-tube is measured as a function of time, first indicating the clot formation and then the clot lysis. For this purpose, we designed a completely computerised semi-automated 8-channel measurement and determination of fibrin clot lysis (EREM, Belgium) (16). Biefly, a computer records every one minute the data from each channel. A software generates the graph of the clot formation and fibrinolytic process. The design of a lysis curve is exemplified in Figure 1A. The *x*-axis/*y*-axis respectively represent time and evolution of the signal sensor. A bloc is made of pure aluminium (360 *×* 30 *×* 100 mm) and is warmed by two resistances of 20 W. It is designed to insert spectrophotometric micro-cuvets (10*×*4*×*45 mm Sarstedt®) into 8 wells. Each well includes one emitter (diode: SFH 409) and one receptor (phototransistor: SFH309FA), both operating at 890 nm. When the clot is formed, the phototransistor signal decreases reflecting a Tyndall effect. For clot formation and lysis analysis, we used purified proteins, t-PA (human recombinant, Hyphen Biomed, Paris, France), PAI-1 (human recombinant, Sigma-Aldrich, Germany), Plasminogen (human plasma purified protein, Sigma-Aldrich, Germany) and Fibrinogen (human plasma purified protein, Sigma-Aldrich). Experimental conditions are presented in the table I.

**TABLE I.** Summary table of the different experimental conditions used for analysis of experimental data and simulations

| Fibrinogen ($mg.ml^{-1}$) | tPA ($\mu g.ml^{-1}$) | PAI-1 ($\mu g.ml^{-1}$) | Pg ($\mu g.ml^{-1}$) | Nb exp. |
| --- | --- | --- | --- | --- |
| 3 | 0.0005 | 0.01 | 13 | 4 |
| 3 | 0.001 | 0.01 | 13 | 18 |
| 3 | 0.002 | 0.01 | 13 | 4 |
| 3 | 0.004 | 0.01 | 13 | 4 |
| 3 | 0.008 | 0.01 | 13 | 4 |
| 3 | 0.001 | 0.0025 | 13 | 4 |
| 3 | 0.001 | 0.005 | 13 | 4 |
| 3 | 0.001 | 0.02 | 13 | 4 |
| 3 | 0.001 | 0.04 | 13 | 2 |
| 3 | 0.001 | 0.01 | 2.6 | 3 |
| 3 | 0.001 | 0.01 | 5.2 | 4 |
| 3 | 0.001 | 0.01 | 26 | 4 |
| 3 | 0.001 | 0.01 | 52 | 4 |
| 0.8 | 0.001 | 0.01 | 13 | 2 |
| 1 | 0.001 | 0.01 | 13 | 2 |
| 2 | 0.001 | 0.01 | 13 | 2 |
| 4 | 0.001 | 0.01 | 13 | 2 |
| 5 | 0.001 | 0.01 | 13 | 1 |
| 6 | 0.001 | 0.01 | 13 | 2 |
|  |  |  |  | total = 74 |

**FIG. 1.**
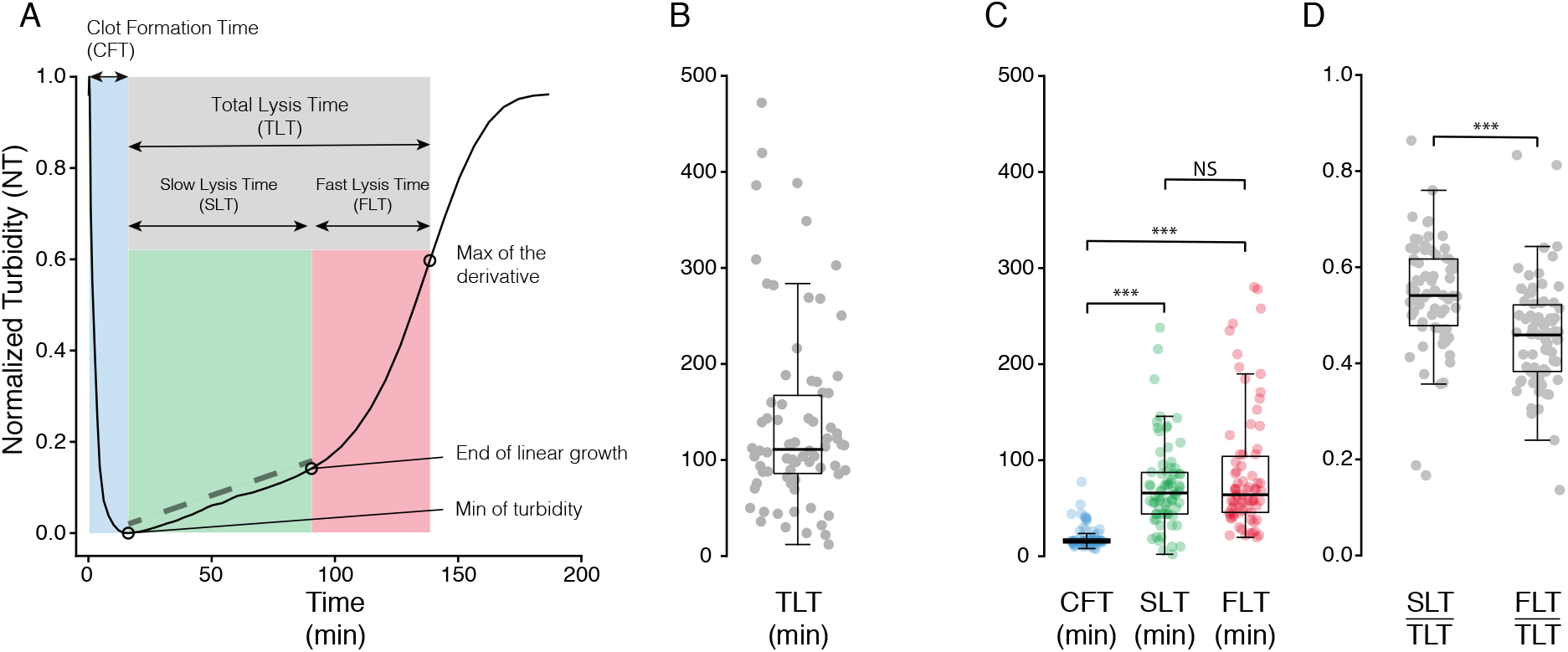
Typical experiment of fibrinolysis and time scales of the processes. A) Turbimetry curve of a fibrinolysis experiment showing the formation of the clot (blue), the total fibrinolysis (gray) and the slow (green) and fast (red) regimes of fibrinolysis. B) Distribution of total lysis time (TLT) over all experiments. C). Distribution of clot formation time (CFT), slow lysis time (SLT), fast lysis time (FLT) over all experiments. D) Ratio of the slow and fast lysis regimes. The sysmbol *∗∗∗* indicates p-values *<* 10^*-*4^.

List of the proteins:

- t-PA (human recombinant, Hyphen Biomed, Paris, France)
- PAI-1 (human recombinant, Sigma-Aldrich, Germany)
- Plasminohen (human plasma purified protein, Sigma-Aldric, Germany)
- Fibrinogen (human plasma purified protein, Sigma-Aldrich)

### Post-treatment and normalization of experimental data

To transform the original turbidity curve to normalized turbidity (NT) we proceeded as follow: The experimental data were post-treated as follow:

1. Original data were sliced and we retained one time step over three.
2. Sliced data were smoothed using Savitzky-Golay filter (from Signal processing scipy.signal library) with third order polynomial fit.
3. Smoothed data were normalized by the maximal value of the turbididy after substracting the minimal value.

We determined the slow lysis regime by fitting the NT curve between the two time points corresponding to NT values of 0.01 and 0.075. Linear fitting was done using with the Optimization and root finding from scipy.optimize. Then to determine the ending point of the linear regime, we identified the time point on the NT curve that is distant from the linear fit by a threshold value 0.005.

### Mathematical models of clot formation and clot lysis

The two models of clot formation and clot lysis are described in details in the main text (see Results and Discussion). We briefly present in this section their main features. The model of clot formation consists in a set of differential equations describing the conversion of fibrinogen to fibrin monomer, the assembly of fibrin monomers into oligomers and the association of protofibrils to form fibrin fibers. The kinetic parameters for the polymerization of fibrin monomers are given Table II and our strategy to estimate their values is detailed in the next section.

**TABLE II.** Simulation parameters for the model of clot formation

| Kinetic parameter | Definition | Value | Unit |
| --- | --- | --- | --- |
| $k_a$ | FPA cleavage | 0.02 | $s^{-1}$ |
| $k_{pi}$ | protofibril initiation | $10^{-17}$ | $L. (molecule\ s)^{-1}$ |
| $k_{pg}$ | protofibril growth | $10^{-15}$ | $L. (molecule\ s)^{-1}$ |
| $k_{fi}$ | fiber initiation | $10^{-18}$ | $L. (molecule\ s)^{-1}$ |
| $k_{fg}$ | fiber growth | $10^{-16}$ | $L. (molecule\ s)^{-1}$ |
| $k_{fa}$ | fiber aggregation | $10^{-18}$ | $L. (molecule\ s)^{-1}$ |

Along with the formation of fibers, PAI-1 can inactivate tPA, Plasminogen and tPA can reversibly bind on the fibers and be activated once they are bound. The kinetic parameters for these reactions are given in Table III.

**TABLE III.** Simulation parameters for the binding and activations of proteins

|  | Kinetic parameter | Value | Unit |
| --- | --- | --- | --- |
| Reversible binding |  |  |  |
| (lys) Plasminogen | $k_f^{Pg}$ | $1.087 \cdot 10^{-4}$ | $\mu M^{-1} s^{-1}$ |
| | $k_r^{Pg}$ | $5.435 \cdot 10^{-5}$ | $s^{-1}$ |
| tPA | $k_f^{tPA}$ | $1.148 \cdot 10^{-4}$ | $\mu M^{-1} s^{-1}$ |
| | $k_r^{tPA}$ | $6.658 \cdot 10^{-5}$ | $s^{-1}$ |
| Acivation of bound species |  |  |  |
| (lys) Plasminogen<sup>b</sup> | $k_{tPA-Pg}^2$ | 0.2 | $s^{-1}$ |
| & tPA<sup>b</sup> | $K_{tPA-Pg}^M$ | 0.02 | $\mu M$ |
| Plasmin<sup>b</sup> | $k_{Pn}^2$ | 25 | $s^{-1}$ |
| | $K_{Pn}^M$ | 250 | $\mu M$ |
| Inactivation |  |  |  |
| tPA<br>& PAI | $k_i^{tPA-PAI}$ | 25 | $\mu M^{-1} s^{-1}$ |

After the formation of the clot, the binding and the activation of the proteins continue (with the same parameters as in Table III) and the lysis starts. The instantaneous rate of lysis takes into the solubilization of fibrin (solubilizaton factor *γ* = 0.1) and is described by a Michaelis-Menten equation.

### Determination of the kinetics parameters for the model of clot formation

We identified 92 sets of plausible parameters that allowed us to reproduce fiber diameters consistent with those measured experimentally. To select the parameters that will be used in our simulations we proceeded as follow:

1. For each experimental conditions we ran the model of fibrinolysis for all the set of parameters
2. For each simulated experiments, we stored the maximal error between experimental and simulated lysis profiles up to the total lysis time.
3. We sorted by increasing values the stored error (each error correspond to a set of parameters) and then scored each set of parameters by summing across experiments the rank of each parameters.
4. We finally selected the set of parameters that had the smallest score. We chose a rank based score to stronger penalize the simulations with the highest errors, no matter the value of the errors.

Considering that we had 92 set of parameters and 74 experiments the worst possible score was 6808, assuming that always the same set of parameters was ranked last. We normalized the rank based score by this value and found that our best solution had a score of 0.22, meaning that on average this set of parameters was in the best quarter. We also found other possible candidates, in particular there were in total 19 set of parameters that were on average in the third of the best solutions (Figure S4). The parameters we retained for the formation of the clot are: and the parameters for reversible binding and activation of proteins are:

## RESULTS AND DISCUSSION

### Analysis of fibrin clot formation and lysis times

Fibrinogen is the key component involved in the blood clotting system. Upon the presence of thrombin, fibrinogen is converted to fibrin monomers after successive cleavages of the fibrinopeptides A and B. Fibrin monomers then self-assemble and polymerize into oligomers which lengthen to form protofibrils and eventually associate to build fibers. Ultimately these fibers branch to form the three-dimensional network thus constituting the clot.

Fibrin fibers are the main substrate for the bindings of the proteins of the fibrinolytic system. Without fibrin, affinity between tissue Plasminogen Activator (tPA) and plasminogen is extremely low and catalytic activation of plasminogen inefficient. However both affinity and efficiency significantly increase in the presence of fibrin, hence favoring the formation of plasmin from the conversion of plasminogen by tPA. On the other hand, fibrinolysis is down regulated by (1) the inhibition of tPA due to the plasminogen activator inhibitor type 1 (PAI-1) and (2) the inhibition of plasmin by the *α*2-antiplasmin.

We analyzed clotting and fibrinolysis using the experimental procedure previously developed in [17] (see Methods), a semi-automatic method that allows one to monitor the dynamics of fibrin assembly and dissolution from turbidity measurements. The actual measurement is the intensity of the light through the system and high turbidity is to be understood as low attenuation, or low opacity. A typical readout is presented in Figure 1A showing the entire time course of an experiment from the formation of the clot (decrease of the turbidity index) to the dissolution of the fibers (increase of the turbidity index). Each step could be identified from the turbidity curve: the clot formation time (CFT) corresponds to the time needed to reach the minimum of turbidity; the total clot lysis time (TLT) is defined as the time interval from the end of the clot formation to the point at which the derivative of the turbidity curve reaches its maximum value (fastest lysis rate). In Figure 1A, the curves were normalized in order to allow comparison between the different experimental conditions (see Methods).

### Experimental Data

Across all our experimental conditions we measured a mean CFT of 19 minutes but the values are broadly distributed (SD of *±*12 minutes), thus indicating that experimental conditions affect the formation time of the clot. We measured a mean TLT of 140 minutes also with a high dispersion (*±*98 minutes SD). Interestingly, we observed that the lysis part of the curve is composed of two regimes. The first one is characterized by a slow increase of turbidity, a slow lysis, denoted SL. The second regime corresponds to a fast increase of turbidity (denoted FL for fast lysis). Independently of the experimental conditions the SL regime appears to be a linear function of time whereas in the FL regime the turbidity increases much faster (Figure S1). It is thus worth discriminating those two processes in the measurement of the TLT. The average duration of the SL regime is around 73 minutes (*±*44 minutes SD). The linear SL regime is characterized by its slope. We found that the inverse of this slope is proportional to the duration of the SL regime (Figure 2A). Because the data are normalized, this result implies that the SL regime might end for a specific value of normalized turbidity (NT). Indeed, the distribution of the values of NT at the end of the SL regime are peaked around 0.1, suggesting that duration of the SL regime might be an intrinsic property (Figure 2B). The FL regime is on average of 67 minutes (*±*56 SD) and comparing the relative contributions of the SL and FL regimes to the TLT, the SL regime accounts for a significantly longer duration than the FL regime (Figure 1D). Altogether these results suggest that over a significant amount of time plasmin-mediated fibrin dissolution is extremely inefficient. Thus, a long slow regime also means a long fast regime. Note that due to the proportionality between the slope of the linear increase of turbidity in the slow regime and the duration of both regimes, it become possible to provide an estimation of the TLT from early measurements (median time 66 min) without waiting hours till the actual end of the lysis. In what follows, the SL regime will also be referred to as the latency regime, due to its ineffective lysis rate.

**FIG. 2.**
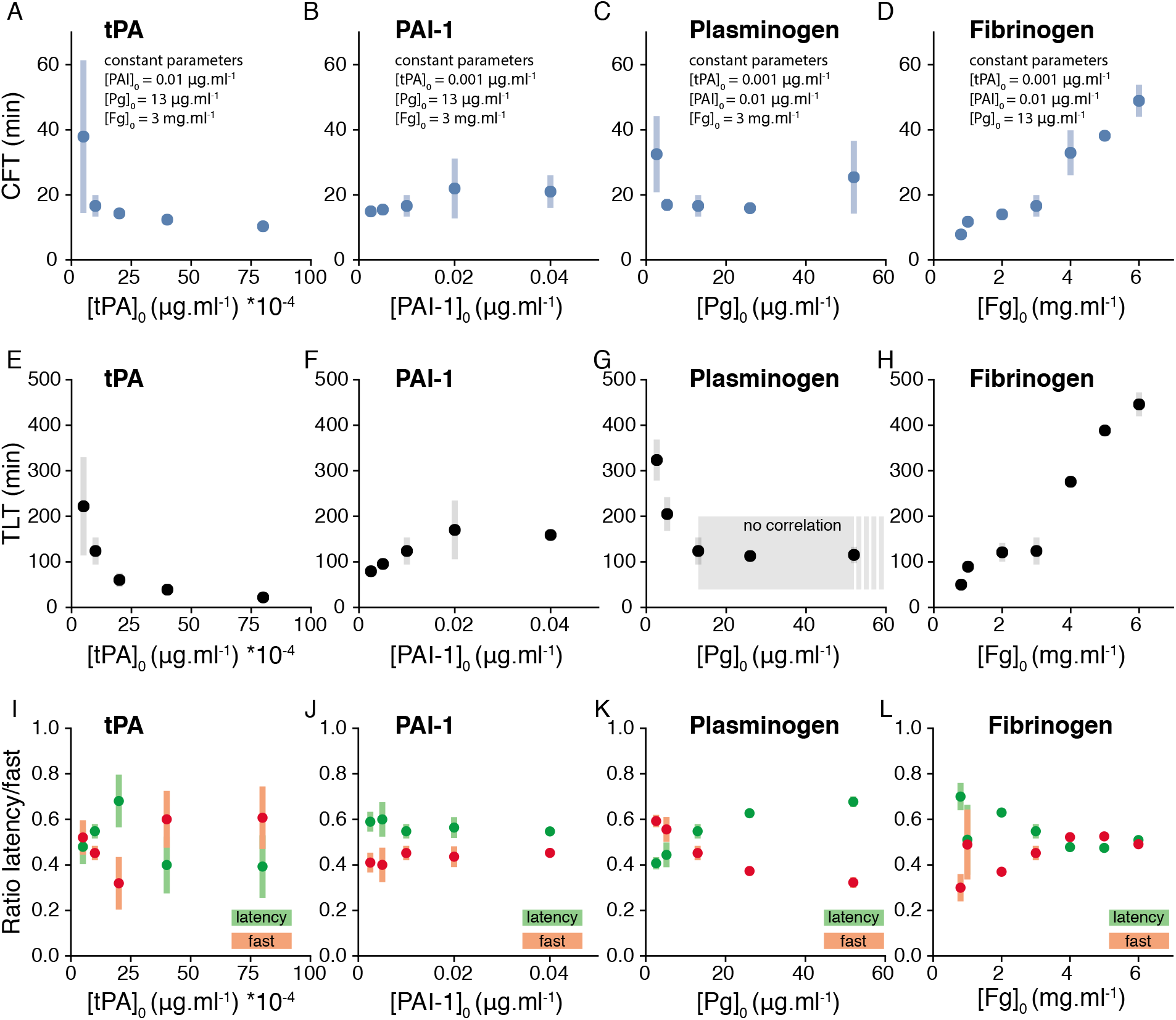
Formation and the lysis time of the clot for different experimental conditions. A-D) Clot formation time (CFT). E-H) Total lysis time (TLT). I-L) Ratio of latency (green) and fast (red) times with respect to the total lysis time. One experimental conditions was varied at a time and all other remained constant. Constant parameters were displayed when relevant.

We then investigated how each event depends on the initial experimental conditions. We compared the clotting time for different concentrations of tPA, PAI-1, plasminogen and fibrinogen (Figure 2A-D). As expected, we found that the initial concentration of tPA has a strong effect on the CFT, the latter decreases as the tPA concentration increases. An explanation is that an excess of tPA rapidly increases plasmin production in the presence of a small amount of fibrin, and it is well known that plasmin can induce the production of a clottable form of fibrinogen [18], thus reducing the time to form the clot.

Conversely the CFT significantly increases with PAI-1 (Spearman correlation *ρ*_*s*_ = 0.44, *p*_*val*_ = 0.016), indicating that PAI-1 inhibits tPA prior to lysis. Considering the time required to form the clot and the fast kinetics of inhibition of tPA by PAI-1 (second order rate constant 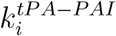 in the order of 10^7^ *M* ^−1^ *s*^−1^ [19]), tPA might bind early during the polymerization of fibrin and the formation of the fibers. If not, most of the initial tPA would be already inhibited and no lysis would be observed. We found no correlation between the CFT and the concentration of plasminogen (*ρ*_*s*_ = 0.05, *p*_*val*_ = 0.853). Instead, the CFT correlates with the initial concentration of fibrinogen mainly because there is more material available to build the clot, as shown by the variation of turbidity Δ*T* between the beginning of the experiment and the end of the clotting (Figure 3). To a lesser extent, Δ*T* decreases (resp. increases) with tPA (resp. PAI-1) jointly with the time needed to form the clot.

**FIG. 3.**
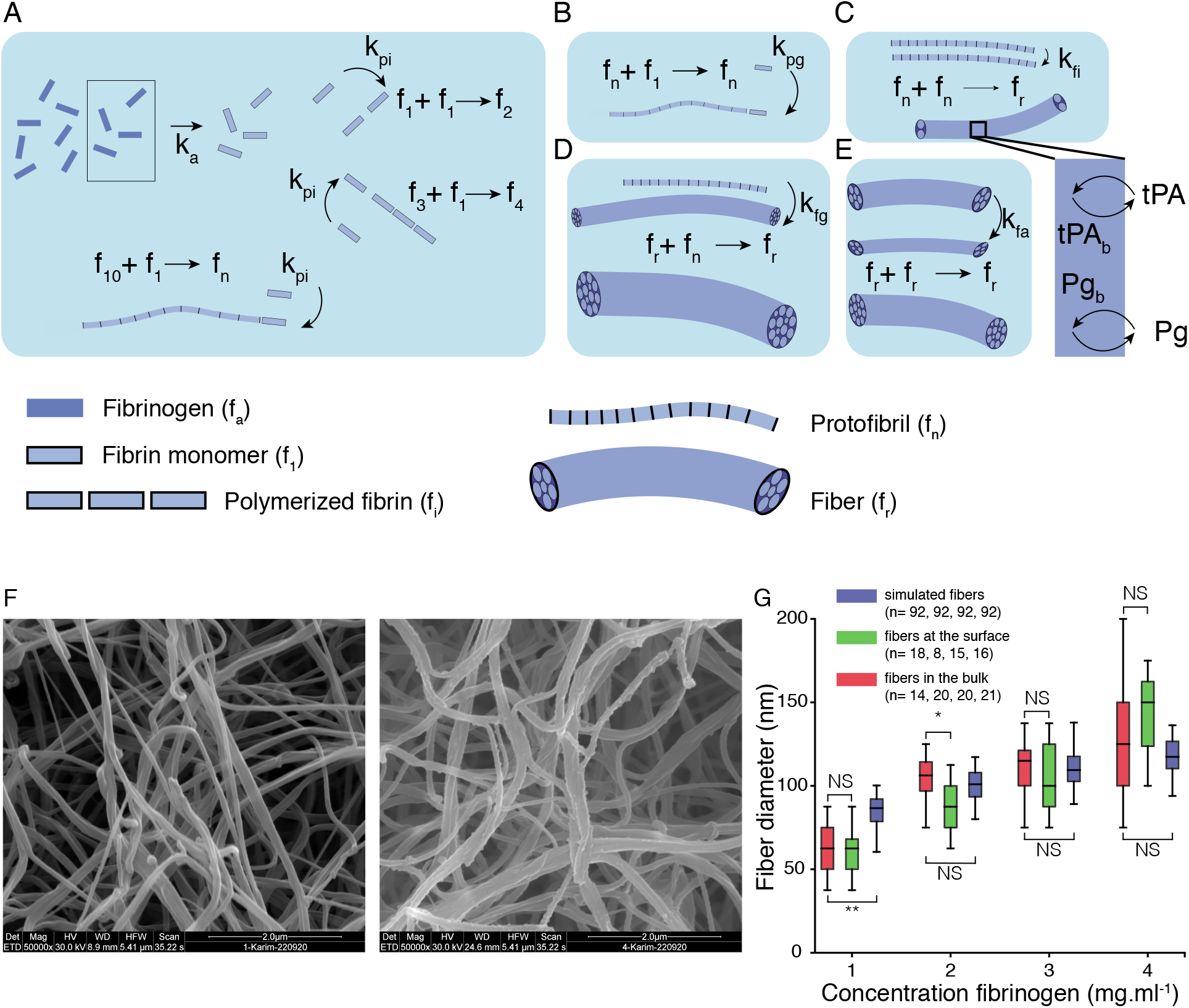
Graphical representation of the model of clot formation. A) Conversion of fibrinogen to fibrin and polymerization of fibrin monomers until the threshold size is reached to form protofibril. B) Protobibrils grow in length by addition of fibrin monomers. C) Association of protofibrils to form fibers. Binding of tPA and plasminogen start when fibers are created. D) Fibers growth by addition of protofibrils. E) Two fibers aggregate together. F) Scanning Electron Microscopy (SEM) images of fibrin fibers at fibrinogene concentration of 1*mg*.*ml*^−1^ (left) and 4*mg*.*ml*^−1^ (right), the scale bar is 2*µm*. G) Boxplot of the fiber diameters for different concentration of fibrinogen measured in the bulk of the clot (red), at the surface of the clot (green) and for simulated fibers (blue).

Then, we investigated the dynamics of the fibrinolysis. As expected from the respective roles of the different components of the fibrinolytic system, the TLT increases with the concentrations of PAI-1 and fibrinogen (Figure 2F and H), and it decreases with the concentration of tPA and Plasminogen. However the effect of plasminogen was mainly significant for concentrations lower than 13*µg*.*ml*^−1^. Comparing side-by-side the SL (or latency) and the FL regimes we could observed how the initial conditions affect the relative contribution of each regime on the total fibrinolysis. For each experiment, we measured the ratios 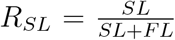 and 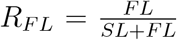 and averaged them over similar experimental conditions. When one of the two ratios is greater than 0.5, the corresponding regime contributes more to the TLT. Regarding the effect of tPA, it appears that the FL regime lasts longer than the SL regime for high initial concentrations of tPA (Figure 2I). For any values of [*PAI -* 1]_0_the SL regime always dominates the fibrinolysis time, showing the importance of the linear regime in the process.

Plasminogen and fibrinogen display opposite trends. Increasing [*Pg*]_0_ corresponds to a decrease of *R*_*F L*_ with a crossing point around [*Pg*]_0_ = 13*µg*.*ml*, whereas an increase of [*Fg*]_0_ corresponds to a increase of *R*_*F L*_. The two curves collapse around [*Fg*]_0_ = 4*µg*.*ml*.

## MODELING CLOT FORMATION AND FIBRINOLYSIS

Our analysis of turbidity curves revealed two important features of the clot formation and lysis process. First the binding of proteins has to be considered during the assembly of the clot, and, second, a significant part of the lysis appears to be inefficient. For the sake of simplicity we decided to decouple the formation of the clot from its lysis and used two different models to describe our experiments. That is to say, we made the assumption that the lysis started once the clot was formed. We adapted a kinetic model of fibrin polymerization from [20] to estimate the concentrations of bound species after clotting and modified the fibrinolysis model from [21] to describe the slow and fast lysis regimes. We hypothesize that the slow and fast regimes are the result of a surface and a bulk binding of the proteins.

### Mathematical model of thrombosis

Formation of a fibrin clot is a step-wise process resulting from the conversion of fibrinogen to fibrin by the cleavage of fibrinopeptide A by thrombin, the polymerization of fibrin monomers to protofibrils and the aggregation of protofibrils to form fibers that eventually merge and branch to create a mesh of fibers (Figure 3). In [20] the authors proposed that protofibrils must reach a minimal length (here an arbitrary length of 11 monomers) before aggregation to account for the lag period observed in turbidity experiments. The equations of the kinetics of the fibers aggregation are the following:

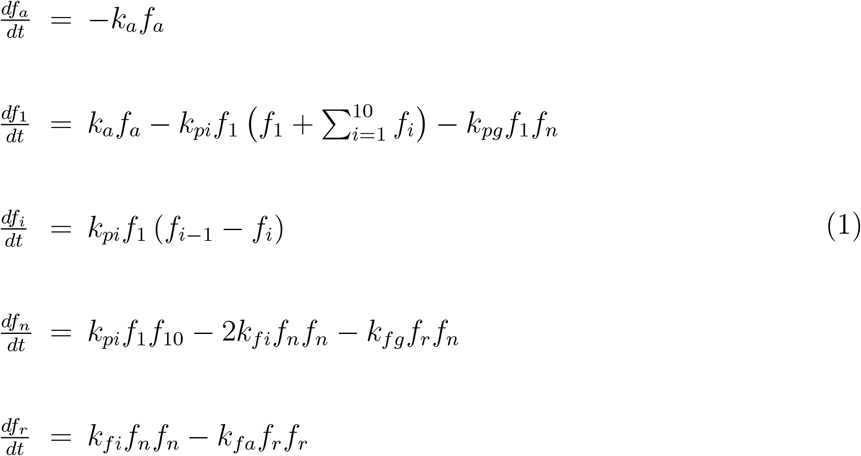

where *f*_*a*_ is the fibrinogen, *f*_*i*_ the fibrin oligomers of size *i, f*_*n*_ the protofibrils and *f*_*r*_represented the fibers. The total amount of protofibrils in fibers 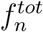, the total amount of fibrin in protofibrils *C*_*fn*_ and the total amount of fibrin in fibers *C*_*fr*_ were given by:

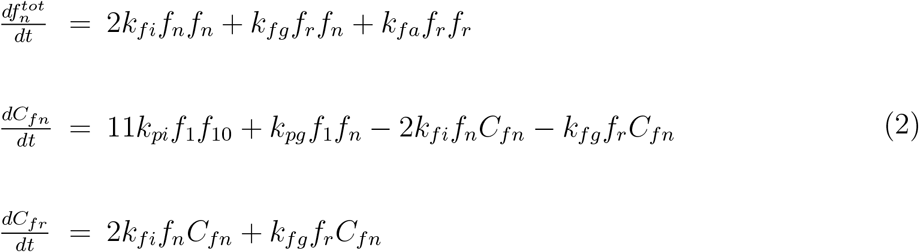

The model described above proposes an over simplified situation of clot formation. For instance it does not account for the three dimensional structure and stability of the fibrin networks which has an impact on the rheological and mechanical properties of the clot as well as on the plasmin-catalyzed fibrinolysis. Whereas more detailed molecular models [22, 23] can predict with greater details fibers properties during network formation, they operate at space and time scales (*nm* and *µs*) that are not compatible with our *in-vitro* experiments (typically hundreds of seconds). The model of Weisel *et al*. proposes a good compromise between molecular and molar levels of description and can be easily extended to determine the amount of binding sites available for reactions during clot formation. Knowing the total number of protofibrils in fibers and the number of fibers, the average fiber radius is given by:

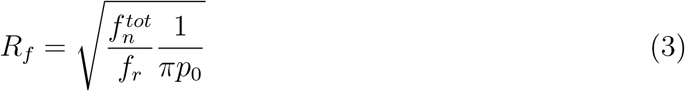

where *p*_0_ = 0.01116 protofibrils/*nm*^2^ is the density of protofibrils in the fiber[24]. Using the radius *r*_0_ of a protofibril [21], it is possible to calculate the number of protofibrils in *κ* layers of a fiber of radius *R*_*f*_ :

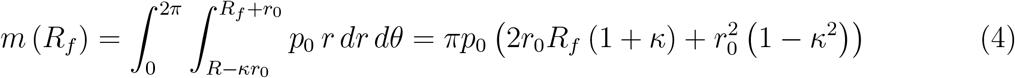

For *κ* = 0, reactions occur at the very surface of the fibers (*i*.*e*. only the exposed side of the outer protofibrils) while for 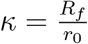 reactions occur in the bulk of the fibers. Using equations 2 we can estimate the number of fibrin per protofibril and, using equation 4, we calculate he number of fibrin on the outer part of the fibers, thus obtaining the number Θ of sites on the fiber available for binding :

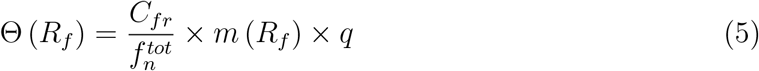

where *q* is the number of binding sites per monomer of fibrin for tPA (*q*_*tPA*_ = 1.5) and plasminogen (*q*_*Pg*_ = 2) [21]. We will discuss later later how changing the values of *κ* Plays a role in the dynamics of the lysis. However during the formation of the fibers we only consider adsorption of the proteins. Knowing the number of binding sites on the fibers during the formation of the clot it is thus possible to estimate the amount of bound and free proteins at the end of the clotting process. Binding was described with second order rate equations, unbinding with first order rate equations and and Michaelis-Menten equation for the activation of plasminogen by bound tPA. The time evolution of these proteins in their free and bound forms are determined with the following set of equations:

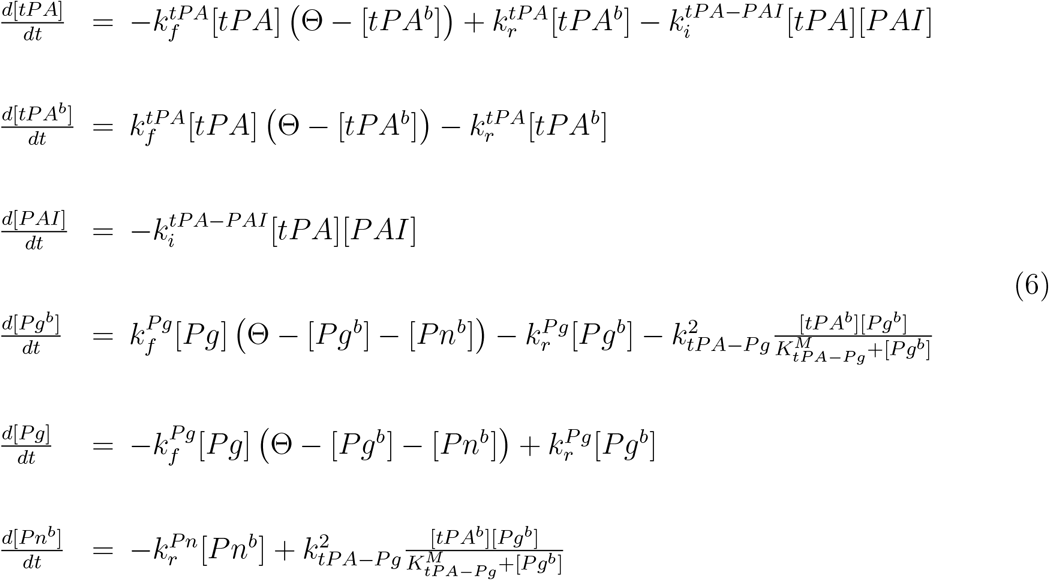

Here we do not account for the direct conversion of plasminogen by tPA in the free phase because it is slow. For the sake of simplicity, we also neglect a potential early lysis simultaneously with the formation of the fibers because the lysis proceeds on time scale much greater than the formation of the clot. The kinetic parameters describing the previous reactions are given in Table III.

The experiments were conducted without flow so does our modeling approach and we do not account for spatial heterogeneity as the different species are homogenized in the initial conditions. The role of the diffusion is discussed elsewhere [25, 26] in the context of sol-gel phase transition.

The experimental values of the kinetic parameters of the polymerization remain largely unknown despite recent experimental progress in live observation of fibrin polymerization [27]. Instead, we identified numerically sets of plausible parameters by solving the equations for a wide range of values *k*_*a*_, *k*_*pi*_, *k*_*pg*_, *k*_*fi*_, *k*_*fg*_, *k*_*fa*_. We ran simulations with initial concentrations of [*tPA*]_0_ = 0.001*µg*.*ml*^−1^, [*Pg*]_0_ = 13*mg*.*ml*^−1^, [*PAI*]_0_ = 0.01*µg*.*ml*^−1^ and concentrations of fibrinogen between 1*mg*.*ml*^−1^ and 4*mg*.*ml*^−1^ with a simulation clotting time of 1000*s* (which was similar to the average CFT for corresponding concentration of fibrinogen). In parallel, we measured fiber diameters using electron microscopy images of clot made in similar experimental conditions (Figure 3F) and we retained kinetics parameters that provide similar fiber diameters (Figure 3G), assuming that at least 80% of the initial fibrinogen is incorporated to the fibers. In total we identified 92 sets of parameters that reproduce the fiber diameters measured experimentally. There is no statistical difference between simulated fiber diameters and fiber diameters measured inside the clot, except at low concentration of fibrinogen, most likely because the simulation time could be too long to converge for these conditions. Among the 92 sets of values, we retained *k*_*a*_ = 0.02*s*^−1^, *k*_*pi*_ = 10^−17^ *L/molecules s, k*_*pg*_ = 10^15^ *L/moleculess, k*_*fi*_ = 10^−18^ *L/moleculess, k*_*fg*_ = 10^−16^ *L/moleculess, k*_*fa*_ = 10^−18^ *L/moleculess* (Table II, Figure S4 and Materials and Methods).

### Mathematical model of fibrinolysis

To model the plasmin-mediated lysis of the fibrin clot we adopted the framework proposed in [21] and more recently used in [28–30]. Briefly, the authors propose that the lysis proceeds from the outside of the fibers, thus reducing their diameter and eventually leading to the lysis of the clot. It is known that plasmin cuts the fibers transversely but it remains debated for a long time whether the fibers were ultimately transected or radially lysed due to uniform distribution of cleavage sites along the fibers. There are now evidences in favor of the first mechanisms both experimentally [31, 32] and theoretically [33, 34]. However in [29] the authors found that transverse lysis and homogeneous shrinkage of the fibers led to similar results.

Knowing the amount of bound plasmin [*Pn*]_*b*_, it is thus possible to estimate the historic amount of lysis *L*(*t*) by integrating over time the instantaneous rate of lysis kinetically described by a Michaelis-Menten equation:

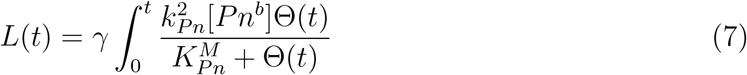

where *γ* = 0.1 is the solubilization factor, *i*.*e* the quantity of fibrin that returns to the plasma phase per number of plasmin cleavage. The percentage of lysis is then obtained by dividing the historical amount of lysis *L*(*t*) by the initial amount of fibrin. Because the lysis proceeds from the outside, the radius of the fiber is directly proportional to the amount of lysis *L*(*t*):

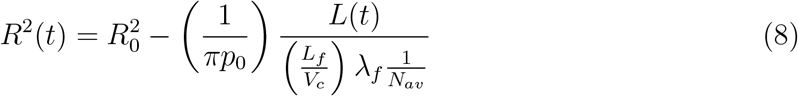

where *L*_*f*_, the average length of the fibers that remains constant during the lysis, could be estimated from geometrical and conservation considerations, *V*_*c*_ is the volume of the clot, *N*_*av*_ is the Avogadro number and *λ*_*f*_ the linear density of fibrin monomers along the fibers. Simultaneously with the decrease of the radius of the fiber, the clot porosity *ϵ* also decreases as:

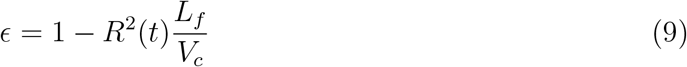

It is important to note that for the fibrinolysis the bound concentration is defined with respect fibrin phase volume. For any specie *i* with an initial concentration in the fluid phase [*i*]_0_ the mass conservation implies that 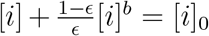. The rates of change for the species in the fluid phase is:

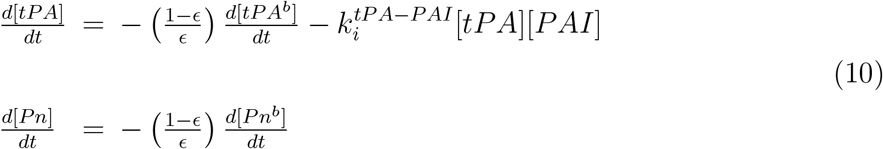

In [21] the authors mainly consider reactions at the outer shell of the fibers but also compare the time to achieve 50% average lysis with different values of *κ*. Nonetheless, with or without the steric constraint they keep the value of *κ* constant during their simulations. Instead, in our work we hypothesize that the number of shells changes dynamically during the lysis, thus explaining the different regimes observed in our experiments. We do not model how the number of shells vary during the lysis, we rather estimate at each time step the optimal value of *κ* that best fits our experimental turbidity curves, based on the fact that the normalized turbidity is a good proxy for the concentration of fibrinogen in the fibers.

In practice we interpolate linearly the turbidity curves between the experimental time points to achieve a similar resolution in time as in simulations (Δ*t* = 0.1*s*). Then at each time step we solve the equations for the fibrinolysis for all possible values of *κ* given the initial conditions provided by the clot formation model (concentrations of bound/free proteins, fibrin incorporated to fibers and radius of the fibers). We estimate the optimal value *κ*^*∗*^ (*t*) as the value of *κ* that at a time *t* minimizes the squared error between *L*^*sim*^ (*t* + Δ*t*), the simulated amount of lysis at the next iteration and the experimental one, given by the normalized turbidity, *NT* (*t* + Δ*t*). Once *κ*^*∗*^ (*t*) is obtained, the equations for the fibrinolysis are actually computed for the time *t* + Δ*t* and lysis continues by repeating the optimization procedure until all fibrin is consumed. Note that the value of *κ*^*∗*^ is estimated based on the time evolution of lysed fibrin and the concentrations of proteins. Thus the difference between *L*^*sim*^ (*t* + Δ*t* and *NT* (*t* + Δ*t*) cannot be made arbitrary small, and errors could potentially accumulate during our fitting procedure. However, we observe that the numerical lysis curves and the turbidity curves do not diverge. This strong consistency between experiment and theory, based on a single parameter *κ* driving the transition from a surface to a bulk reaction, reinforces the credibility of our model.

It is worth recalling that in our approach the formation of the clot and its lysis do not occur at the same time. Instead, the two models are coupled assuming a time scale separation. For each simulation, we determine the initial conditions for the lysis from the output of the clot formation module. As a validation of the lysis model, we compare directly the simulated and its corresponding experimental lysis profiles. Our results are summarized in Figure 4. The change in the experimental conditions are indicated on each figure, while all other concentrations stay the same. Overall we find a very good agreement between the experiments and the simulations, both being most of the time indistinguishable. In particular the results for the change of tPA match almost perfectly the experiments (Figure4A), which is of great interest considering the fact that a critical component of the clinical treatment is the amount of tPA injected to the patients.

**FIG. 4.**
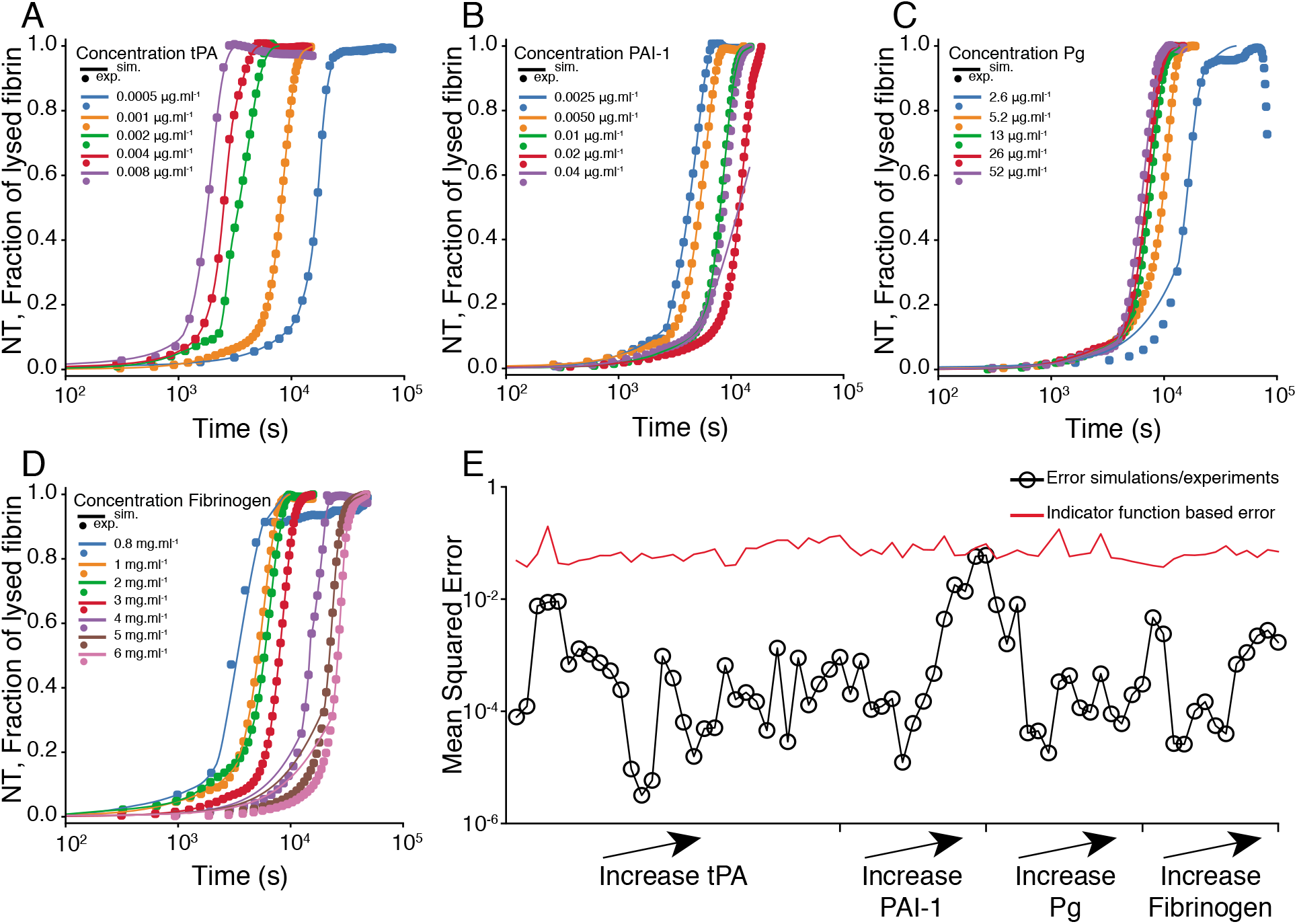
Comparison of simulated and experimental lysis profiles. Time course of normalized turbidity (NT) and fraction of lysed fibrin in simulated for different initial concentrations of A) tPA, B) PAI-1, C) Plasminogen (Pg), D) Fibrinogen, E) Mean squared error for each simulation (black) and mean square error for a surrogate lysis profile (red) described by an indicator function 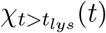, where *t*_*lys*_ is the lysis time.

Note that our model poorly captures the experimental FL regime for clots produced at a high PAI-1 concentration. Numerical simulations predict a long TLT, as one would expect considering that PAI-1 inhibits tPA. Yet, the TLT is experimentally very short (Figure 2). We cannot exclude an experimental bias in this case: two out of the four experiments have a short TLT while the other ones have no lysis at all (data not shown).

Another discrepancy between simulation and experiment is observed for very high concentrations of fibrinogen (Figure 4D). Between the end of the SL regime and the start of the FL regime, the model fails to reproduce the observations. It is possible that this difference between the simulations and the experiments is due to the initial conditions, either a too high amount of bound tPA, or a too small fiber radius. For all the simulations we estimate the initial conditions from a single set of parameters for the thrombosis. It is possible that the kinetic parameters for the clot formation are different at high concentrations of fibrinogen. For instance, this could be due to fibers interactions, thus changing the estimated amount of bound tPA. Another possibility explaining this difference is that the competition between clot formation and lysis starts earlier than expected. If so, at the estimated starting point of the lysis the clot already begins to dissolve and some bound tPA is already consumed, resulting in an overestimation of the tPA concentration in the initial lysis conditions.

Nonetheless our results are globally very good. To quantify the agreement between simulations and experiments, the average squared error between simulated and experimental lysis curves is measured. We find a minimum error of 3 *×* 10^−6^ and a maximum of 6 *×* 10^−2^ (Figure4E, black curve). In order to understand the meaning of these values, we compare them with the mean squared error that would be obtained if the lysis profiles were described by indicator functions 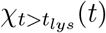 (*t*), where *t*_*lys*_ is the corresponding lysis time.

We then investigate the change over time of the fiber radius (Figure 5). As expected, the radius of the fibers invariably decrease as the lysis proceeds. For all conditions we observe two regimes, the first one in which the radius decreases slowly, and the second one where the radius drops sharply to a value close to 0. This behavior obviously reminds us the the SL and FL regimes illustrated in Figure 1. In Figure 5A-D, we add the experimentally obtained time points corresponding to the end of the SL and Fl regimes on top of the simulation curves giving the evolution of the fiber radius. These time points are displayed as circles and indicate that the slow and fast predicted radius evolution regimes take place at times that are in agreement with the slow and fast lysis regimes observed experimentally.

**FIG. 5.**
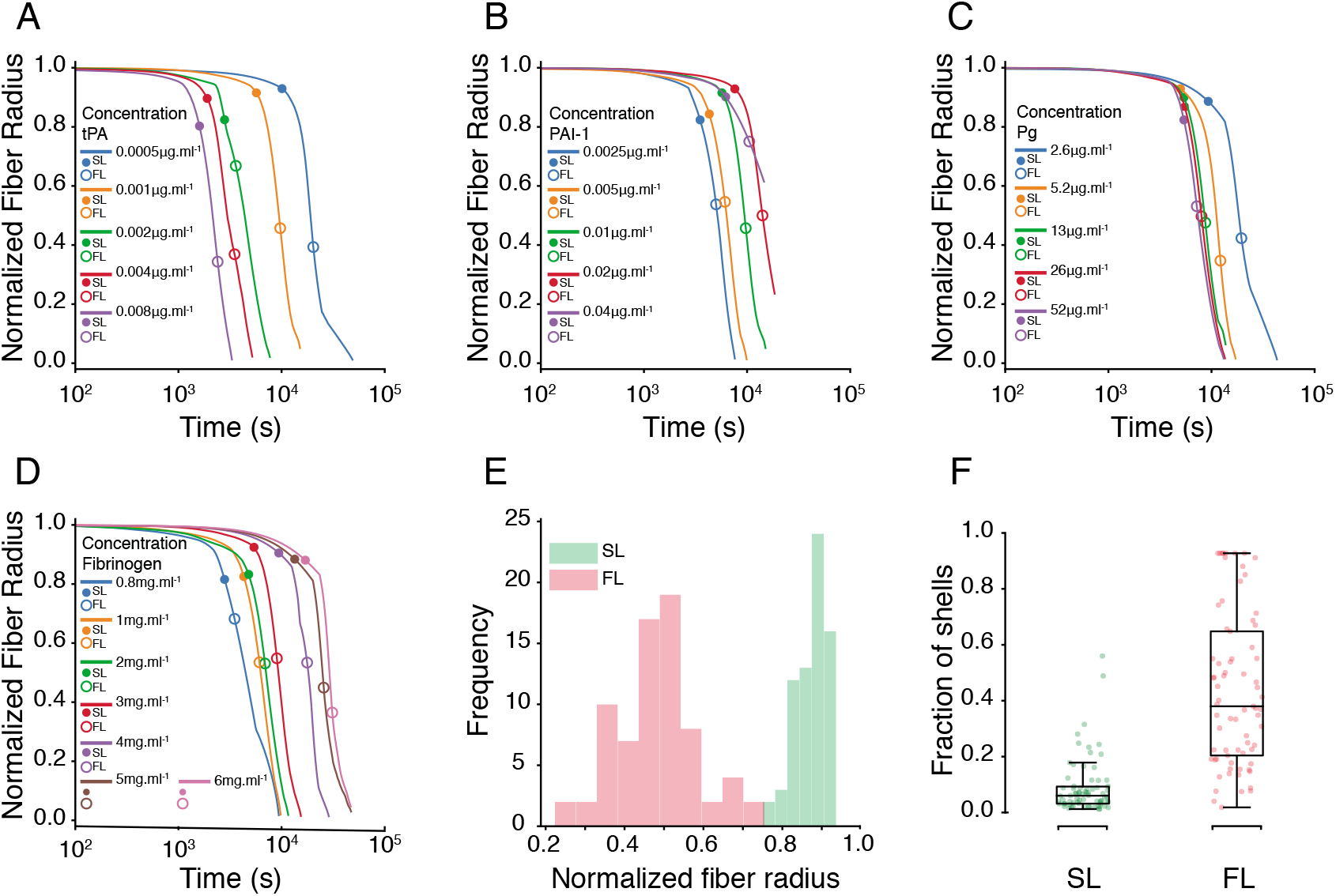
Time evolution of the radius of the fibers during fibrinolysis. Time change of the normalized fiber radius for various concentrations of A) tPA, B) PAI-1, C) Plasminogen (Pg), D) Fibrinogen. Distribution of the values of the normalized fiber radius at the end of the SL regime (green) and FL regime (red). F) Boxplot of the fraction of shells participating to the reactions during the SL regime (green) and the FL regime (red). Green (resp. red) points represent the time average of the fraction of shell during the latency regime (resp. between end of latency and end of fast regime) participating to reactions.

Furthermore, the distributions of the normalized fiber radius at the end of each regime are both nicely peaked, around 0.5 for the FL regime and 0.9 for the SL regime. This result reminds us of the distribution of the values of the normalized turbidity at the end of the SL regime (see Figure S2). The fact that the distributions of fiber radius in both regimes are statistically well separated supports our hypothesis that the two lysis regimes we observed in our *in vitro* experiments can be explained by a change of the number of shells involved in the kinetics of fibrinolysis. As a further indication, we aggregated for all simulations the percentage of shells participating to the bindings of proteins during the SL and FL regimes. During the SL regime around 8% of the shells are involved while in the FL regime this value is around 47%. Consequently, to sustain a regime of slow lysis it is sufficient that only few shells participate to the reactions.

## CONCLUSION

While the interplay between the main components of the fibrinolytic system is well understood, some dynamical aspects of the fibrinolysis remain unclear. Notably, we observe that in vitro fibrin rich clots undergo a slow and inefficient phase of degradation when subjected to endogenous fibrinolysis. Actually a large part of the lysis process operates in this slow regime. To explain this observation, we proposed a computational model in which the properties of the binding of the proteins change during the lysis. First plasminogen and tissue plasminogen activator bind at the surface of the fibers, resulting in a slow lysis, then in the bulk of the fibers thus speeding up the degradation of the clot.

Our model differs from the seminal work of Diamond and Anand [21] and the latest works by Piebalgs, Xu and co-workers [29, 30] because we consider here that the number of binding sites accessible for the adsorption of soluble species dynamically change during the lysis. We hypothesize that this mechanisms allows us to capture the different regimes of lysis observed in our experiments. Identifying such transition from surface to bulk reactions may be difficult to observe experimentally. However following lysis of a fiber in real time, Feller et al [35] discussed a phenomenological fiber-level model of fibrinolysis with different time scales based on the integrity of the fibers. They proposed that plasmin first cleaves inter-protofibrils, loosening fibers and making accessible low-affinity sites on fibers. Then a rapid phase of lysis follows, due to axial fragmentation of the protofibrils. Our hypothesis of dynamical change of binding sites accessibility may implement a macroscopic version of the aforementioned phenomenological mechanism.

Note that in the present study, we neither investigate experimentally the effect of the plasmin inhibitor *α*-2-antiplasmin nor the impact of thrombin-activatable fibrinolysis inhibitor (TAFI) on the lysis time. However the linear regime was also observed in an euglobulin clot lysis assay [36], thus indicating that the linear regime also holds in more complex situation.

## AUTHOR CONTRIBUTIONS

FR, KZB and BC designed the study and wrote the manuscript. FR developed the numerical models and performed data analysis. KZB and AR performed experiments. DM and DPM provided SEM images and performed SEM image analysis. KBZ and BC supervised the project.

## ACKNOWLEDGMENTS

This work was supported by grants from the INSIST project (grant #777072 from the European Commission Horizon2020 program)the CHU Charleroi; the Fonds de la Chirurgie Cardiaque; Fonds de la Recherche Medicale en Hainaut (FRMH).

## SUPPLEMENTARY MATERIAL

**Supplementary Figure 1.**
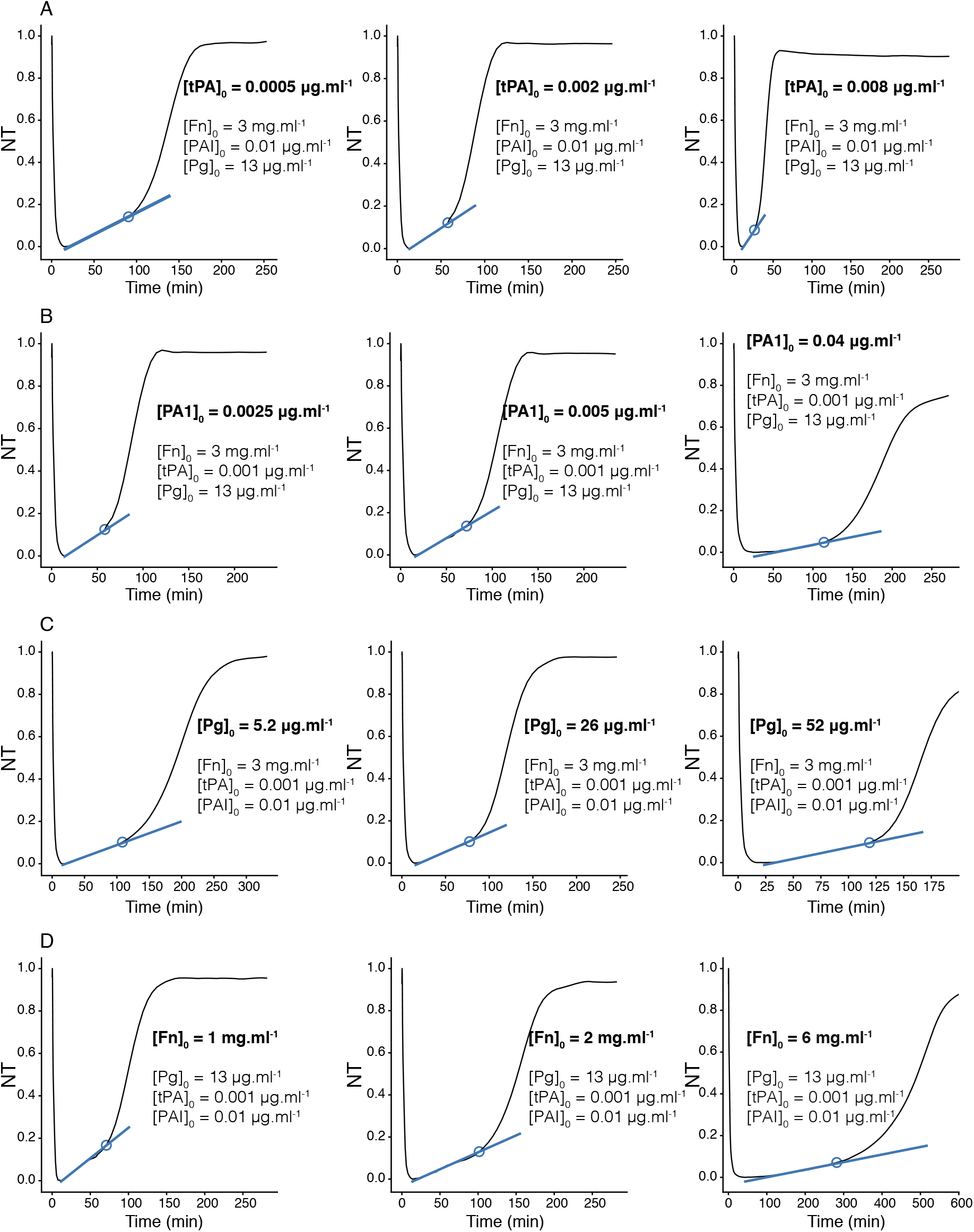
Normalized turbidity (NT) increased linearly in time during the slow regime. Normalized turbidity time curve for different concentrations of A) tPA, B) PAI-1, C) Plasminogen, D) Fibrinogen.

**Supplementary Figure 2.**
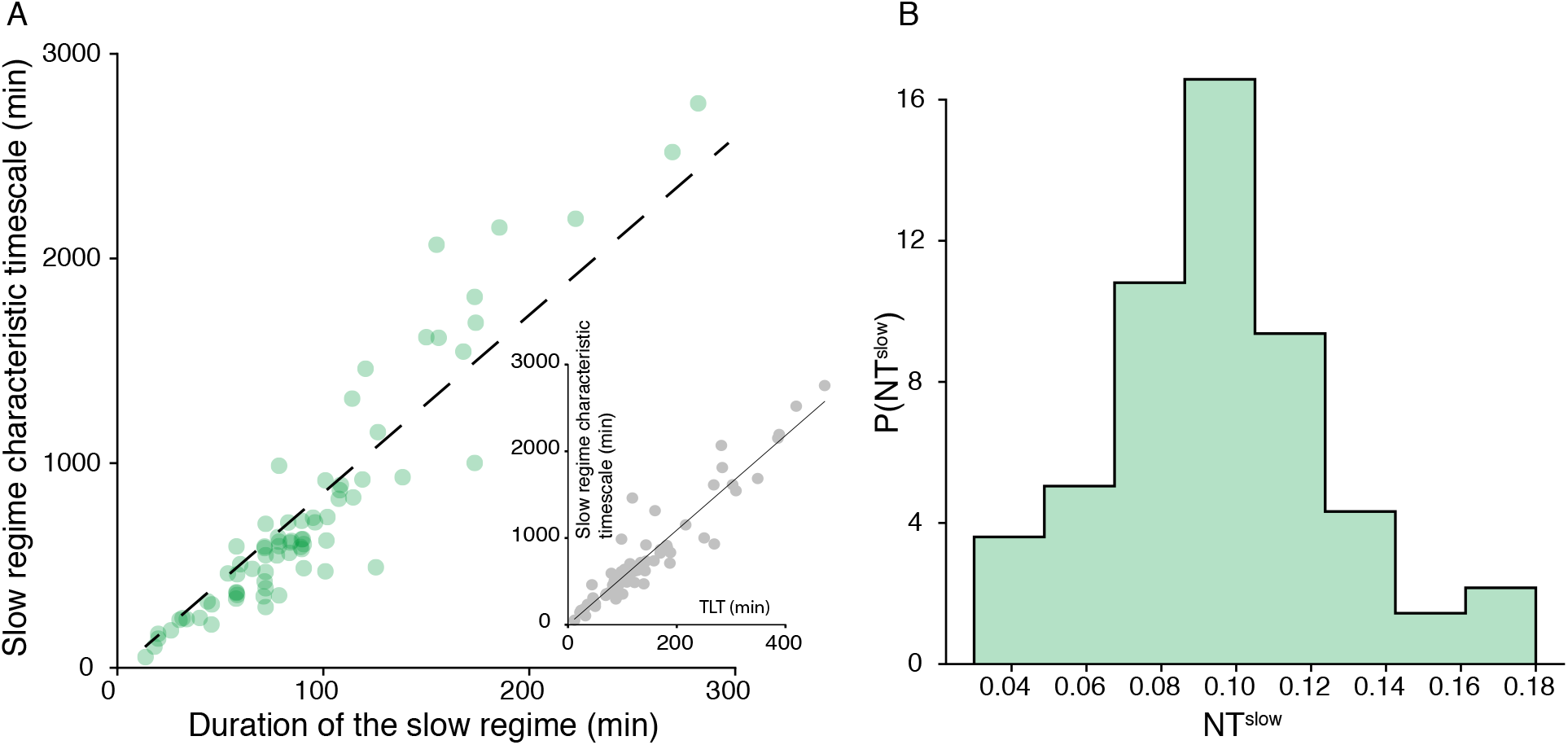
Timescale and duration of the slow regime correlated. A) Inverse of the slope of the linear fit function of the duration of the slow regime (inset: Total lysis time). B) Normalized histogram of the values of the normalized turbidity at the end of the slow regime.

**Supplementary Figure 3.**
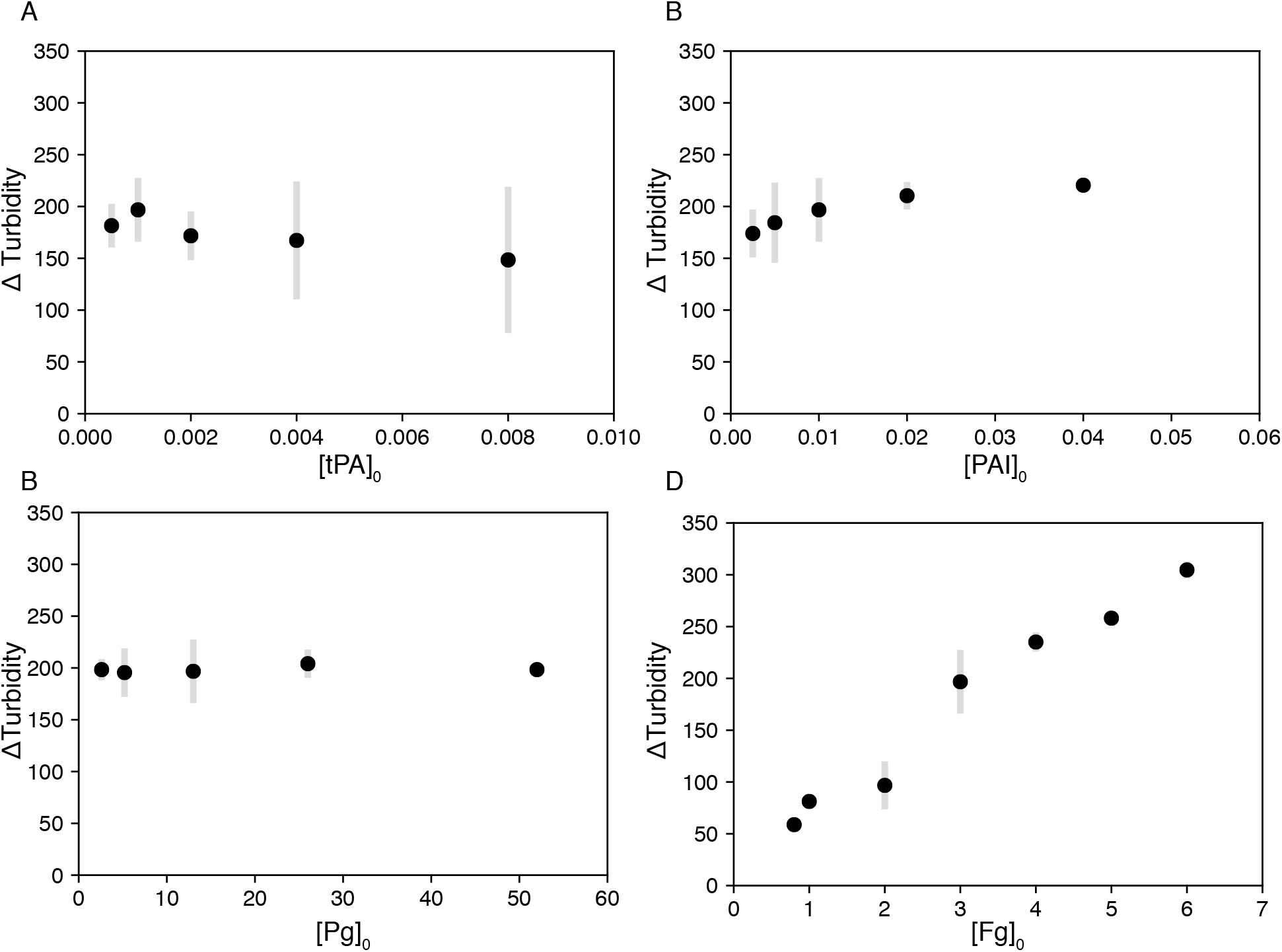
Variation of the difference max to min of turbidity (Δ Turbidity for different concentrations of A) tPA, B) PAI-1, C) Plasminogen, D) Fibrinogen.

**Supplementary Figure 4.**
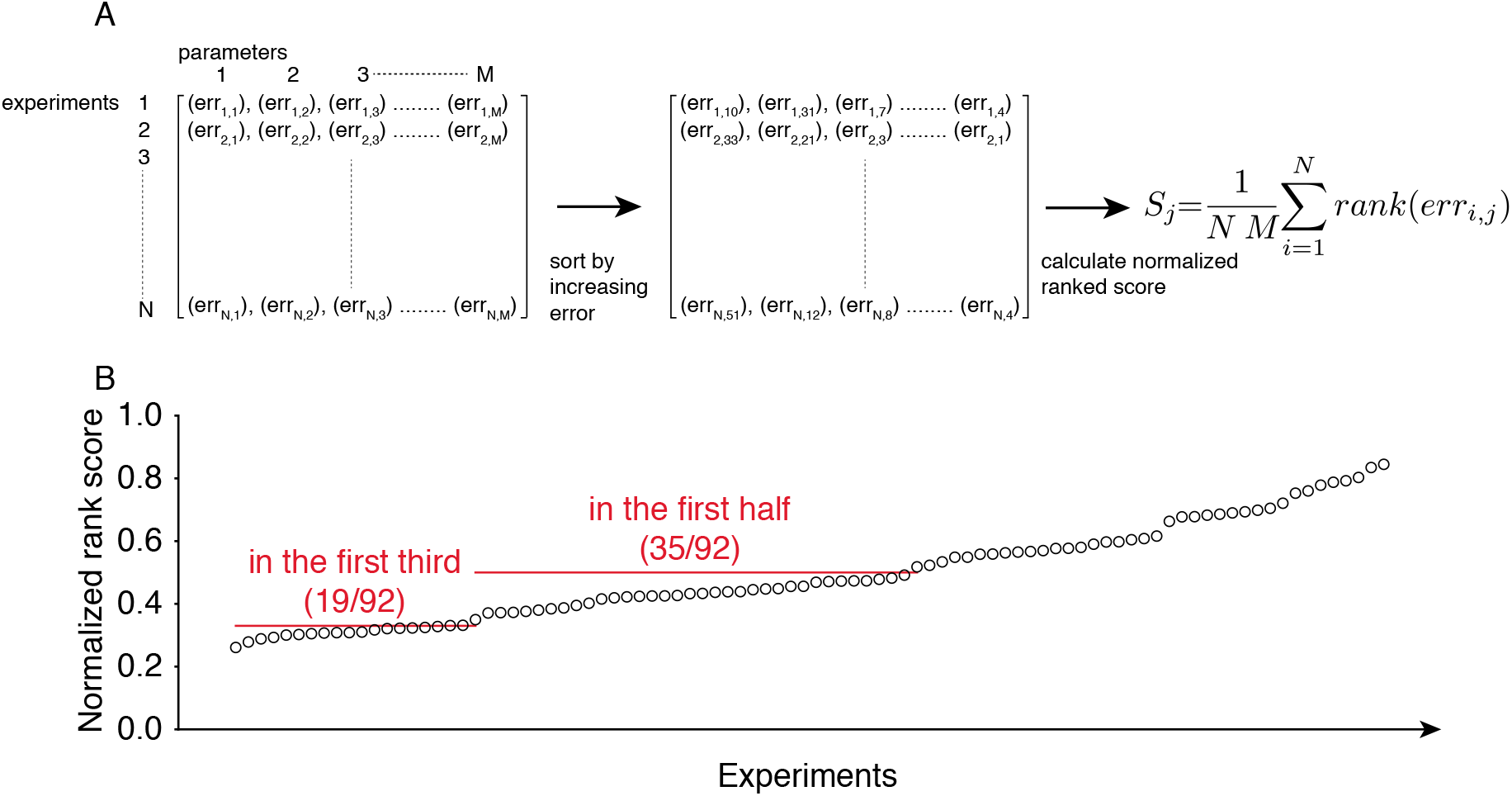
Scoring the kinetic parameters of the clot formation model. A) Methodology to determine the score of each set of kinetic parameters. First we measured err_*ij*_ the maximum error between simulated and experimental lysis profile for each experiment *i* and each set of parameters *j*. Then we sorted the errors for each experimental condition and summed their corresponding rank to calculate the score *S*. B) Distribution of the score for all the simulations.

## References

[1] K. C. Robbins, G. H. Barlow, G. Nguyen, and M. M. Samama, Comparison of plasminogen activators, Seminars in Thrombosis and Hemostasis 13, 131 (1987).

[2] K. Zouaoui Boudjeltia, M. Guillaume, C. Henuzet, P. Delr’ see, P. Cauchie, C. Remacle, J. Ducobu, M. Vanhaeverbeek, and D. Broh’ee, Fibrinolysis and cardiovascular risk factors: As-sociation with fibrinogen, lipids, and monocyte count, European Journal of Internal Medicine 17, 102 (2006).

[3] M. Pahor, M. B. Elam, R. J. Garrison, S. B. Kritchevsky, and W. B. Applegate, Emerging noninvasive biochemical measures to predict cardiovascular risk, Archives of Internal Medicine 159, 237 (1999).

[4] V. Salomaa, V. Stinson, J. D. Kark, A. R. Folsom, C. E. Davis, and K. K. Wu, Association of fibrinolytic parameters with early atherosclerosis: The ARIC study, Circulation 91, 284 (1995).

[5] S. G. Thompson, J. Kienast, S. D. Pyke, F. Haverkate, and J. C. van de Loo, Hemostatic Factors and the Risk of Myocardial Infarction or Sudden Death in Patients with Angina Pectoris, New England Journal of Medicine 332, 635 (1995).

[6] J. D. Pearson, 6 The control of production and release of haemostatic factors in the endothelial cell, Bailliere’s Clinical Haematology 6, 629 (1993).

[7] Top 10 causes of death, https://www.who.int/gho/mortality_burden_disease/causes_death/top_10/en/m, accessed: 2010-09-30.

[8] J. Minnerup, H. Wersching, A. Teuber, J. Wellmann, J. Eyding, R. Weber, G. Reimann, W. Weber, L. U. Krause, T. Kurth, K. Berger, V. Homberg, A. Petrovitch, L. Heuser, P. Mönnigs, C. Krogias, B. Wallner, S. Hennigs, A. Ahlers, H. Sahl, A. Ranft, C. Dobis, F. Brassel, M. Nolden-Koch, H. Schmitt, R. Chapot, H. Nordmeyer, M. Schlamann, C. Weimar, F. Busch, E. W. Busch, E. Sigges, H. Ruf, K. Wohlfahrt, R. Karatschai, B. Klein, T. Höhle, A. Haass, A. Nasreldein, B. Büchele, G. Gahn, M. Sterker, T. Hantel, C. Krämer, H. Henningsen, I. Adelt, M. König, C. Schmidt, A. Hofmann, T. Niederstadt, M. Unrath, T. Rehfeldt, B. Fauser, A. Pfeiffer, S. Lowens, F. Stögbauer, T. Staudacher, P. Erdmann, K. H. Grotemeyer, E. Spüntrup, P. Bücke, P. Wienecke, J. Faiss, M. Wolzik-Großmann, N. Brune, S. Isenmann, C. Thomas, and D. Mucha, Outcome after Thrombectomy and Intravenous Thrombolysis in Patients with Acute Ischemic Stroke: A Prospective Observational Study, Stroke 47, 1584 (2016).

[9] E. A. Mistry, A. M. Mistry, M. O. Nakawah, R. V. Chitale, R. F. James, J. J. Volpi, and M. R. Fusco, Mechanical Thrombectomy Outcomes with and Without Intravenous Thrombolysis in Stroke Patients: A Meta-Analysis, Stroke 48, 2450 (2017).

[10] E. The ATLANTIS and N. rt PA Study Group Investigators, Association of outcome with early stroke treatment: Pooled analysis of ATLANTIS, ECASS, and NINDS rt-PA stroke trials, Lancet 363, 768 (2004).

[11] S. B. Coutts, E. Berge, B. C. Campbell, K. W. Muir, and M. W. Parsons, Tenecteplase for the treatment of acute ischemic stroke: A review of completed and ongoing randomized controlled trials, International Journal of Stroke 13, 885 (2018).

[12] P. R. Konduri, H. A. Marquering, E. E. van Bavel, A. Hoekstra, C. B. L. M. Majoie, and T. I. I., In-silico trials for treatment of acute ischemic stroke, Frontiers in Neurology 11, 1062 (2020).

[13] M. F. Hockin, K. C. Jones, S. J. Everse, and K. G. Mann, A model for the stoichiometric regulation of blood coagulation, Journal of Biological Chemistry 277, 18322 (2002).

[14] M. S. Chatterjee, W. S. Denney, H. Jing, and S. L. Diamond, Systems Biology of Coagulation Initiation: Kinetics of Thrombin Generation in Resting and Activated Human Blood, PLoS Computational Biology 6, e1000950 (2010).

[15] C. M. Danforth, T. Orfeo, K. G. Mann, K. E. Brummel-Ziedins, and S. J. Everse, The impact of uncertainty in a blood coagulation model, Mathematical Medicine and Biology 26, 323 (2009).

[16] A. Shibeko, B. Chopard, A. Hoekstra, and M. Panteleev, Redistribution of TPA fluxes in the presence of PAI-1 regulates spatial thrombolysis, Biophysical Journal 10.1016/j.bpj.2020.06.020 (2020).

[17] K. Z. Boudjeltia, P. Cauchie, C. Remacle, M. Guillaume, D. Broh’ see, J. L. Hubert, and M. Vanhaeverbeek, A new device for measurement of fibrin clot lysis: application to the euglobulin clot lysis time, BMC biotechnology 2, 8 (2002).

[18] G. Cesarman-Maus and K. A. Hajjar, Molecular mechanisms of fibrinolysis, British Journal of Haematology 129, 307 (2005).

[19] M. E. Carr and B. M. Alving, Effect of fibrin structure on plasmin-mediated dissolution of plasma clots, Blood Coagulation and Fibrinolysis 6, 567 (1995).

[20] J. W. Weisel and C. Nagaswami, Computer modeling of fibrin polymerization kinetics correlated with electron microscope and turbidity observations: clot structure and assembly are kinetically controlled, Biophysical Journal 63, 111 (1992).

[21] S. Diamond and S. Anand, Inner clot diffusion and permeation during fibrinolysis, Biophysical Journal 65, 2622 (1993).

[22] S. Yesudasan, X. Wang, and R. D. Averett, Coarse-grained molecular dynamics simulations of fibrin polymerization: effects of thrombin concentration on fibrin clot structure, Journal of Molecular Modeling 24, 109 (2018).

[23] S. Yesudasan, X. Wang, and R. D. Averett, Fibrin polymerization simulation using a reactive dissipative particle dynamics method, Biomechanics and Modeling in Mechanobiology 17, 1389 (2018).

[24] M. E. Carr and J. Hermans, Size and Density of Fibrin Fibers from Turbidity, Macromolecules 11, 46 (1978).

[25] A. Lobanov, A. Nikolaev, and T. Starozhilova, Mathematical Model of Fibrin Polymerization, Mathematical Modelling of Natural Phenomena 6, 55 (2011).

[26] A. I. Lobanov, Fibrin polymerization as a phase transition wave: A mathematical model, Computational Mathematics and Mathematical Physics 56, 1118 (2016).

[27] A. Hategan, K. C. Gersh, D. Safer, and J. W. Weisel, Visualization of the dynamics of fibrin clot growth 1 molecule at a time by total internal reflection fluorescence microscopy, Blood 121, 1455 (2013).

[28] E. Khramchenkov and M. Khramchenkov, Numerical simulation of rheological, chemical and hydromechanical processes of thrombolysis, in Journal of Physics: Conference Series, Vol. 602 (Institute of Physics Publishing, 2015) p. 012042.

[29] A. Piebalgs and X. Y. Xu, Towards a multi-physics modelling framework for thrombolysis under the influence of blood flow, Journal of The Royal Society Interface 10.1098/RSIF.2015.0949 (2015).

[30] A. Piebalgs, B. Gu, D. Roi, K. Lobotesis, S. Thom, and X. Y. Xu, Computational Simulations of Thrombolytic Therapy in Acute Ischaemic Stroke, Scientific Reports 8, 10.1038/s41598-018-34082-7 (2018).

[31] J. P. Collet, D. Park, C. Lesty, J. Soria, C. Soria, G. Montalescot, and J. W. Weisel, Influence of Fibrin Network Conformation and Fibrin Fiber Diameter on Fibrinolysis Speed, Arteriosclerosis, Thrombosis, and Vascular Biology 20, 1354 (2000).

[32] A. Blinc, J. Magdic, J. Fric, and I. Musevic, Atomic force microscopy of fibrin networks and plasma clots during fibrinolysis, Fibrinolysis and Proteolysis 14, 288 (2000).

[33] B. E. Bannish, J. P. Keener, and A. L. Fogelson, Modelling fibrinolysis: a 3D stochastic multiscale model, Mathematical Medicine and Biology: A Journal of the IMA 31, 17 (2014).

[34] B. E. Bannish, I. N. Chernysh, J. P. Keener, A. L. Fogelson, and J. W. Weisel, Molecular and Physical Mechanisms of Fibrinolysis and Thrombolysis from Mathematical Modeling and Experiments, Scientific Reports 7, 10.1038/s41598-017-06383-w (2017).

[35] T. Feller, J. H’ sarsfalvi, C. Cs’anyi, B. Kiss, and M. Kellermayer, Plasmin-driven fibrinolysis in a quasi-two-dimensional nanoscale fibrin matrix, Journal of Structural Biology 203, 10.1016/j.jsb.2018.05.010 (2018).

[36] A. A. Smith, L. J. Jacobson, B. I. Miller, W. E. Hathaway, and M. J. Manco-Johnson, A new euglobulin clot lysis assay for global fibrinolysis, Thrombosis Research 112, 329 (2003).

